# CIRI-hub: an integrated and visual analytics platform for circular RNAs in cancers

**DOI:** 10.1101/2023.08.09.552604

**Authors:** Wanying Wu, Jinyang Zhang, Fangqing Zhao

## Abstract

Recent studies have demonstrated the emerging functions of circular RNAs (circRNAs) in regulating tumor progression and metastasis, and various databases have been established for exploring the expression patterns and features of circRNAs. However, all these databases only provide simple browsing or searching functions with a limited collection of samples, and none of them provides integrated analytical functions of given circRNAs or in-house RNA-Seq data. Here, we developed CIRI-hub, which provides a user-friendly webserver for integrative analyses of circRNAs and outputting publication-quality figures. CIRI-hub integrates a compendium of 2187 tumor and 680 normal RNA-seq libraries spanning 33 tissue types. The CIRI-hub webserver can accept various formats of input, and perform automated analysis of circRNAs in user input data as well as a variety of tumor and matching normal tissues. All analysis results can be downloaded as vectorized figures, and thresholds and plotting parameters can be interactively customized using the visual interface. We believe that CIRI-Hub will serve as a powerful tool for identifying novel cancer biomarkers and exploring the biological functions of circRNAs in tumorigenesis.

**Key Points:**

1. CIRI-hub integrates a compendium of 2187 tumor and 680 normal RNA-seq libraries spanning 33 tissue types, and provides comprehensive analysis and visualization functions for pan-cancer circRNA analysis.
2. CIRI-hub permits users to specific circRNAs of their interests or upload their dataset and provides interactively customization of analysis algorithms and plotting parameters on the web interface.
3. CIRI-hub can serve as a powerful tool for identifying novel cancer biomarkers and exploring the biological functions of circRNAs in tumorigenesis.

## Background

Circular RNAs (circRNAs) are a large class of RNA molecules with covalent circular structures. It has been reported that circRNAs can play important roles in sponging miRNAs [1, 2] and proteins [3, 4], regulating transcription and splicing [3, 5], and translating proteins through internal ribosome entry sites (IRES) and cap-independent manners [6, 7]. In addition, recent studies have demonstrated the highly specific expression of circRNA across various tumor types and revealed the emerging roles of circRNAs in regulating tumor progression and metastasis [8-10]. Previous studies have shown that circRNAs could have specific internal structures like novel exons [11-14], which suggested the unique biological functions of circRNAs. Furthermore, the lack of linear 5’ and 3’ ends makes circRNAs resistant to exonuclease-mediated degradation compared to their linear counterparts [15]. The high stability of circRNAs suggested the presumable possibility of developing circRNAs as promising diagnostic biomarkers or therapeutic targets across different cancer types.

With the advent of sequencing technology, an enormous number of cancer-related data have been generated. In recent years, various databases have been developed for exploring the expression pattern and features of circRNAs in cancers [16-21]. For instance, MiOncoCirc [18] characterized circRNAs across >2,000 cancer samples, and circRic [19] and CSCD2 [20] also provide comprehensive resources for cancer-specific circRNAs in human cancer and cell lines. However, all these databases only focus on browsing and searching of public data and analysis results, and none of these websites provides a convenient portal for customized analysis and visualization of users’ datasets, which is a bottleneck for most web lab users.

To fill this gap, we present CIRI-hub, a web server for automated analysis of circRNAs in tumors and matching normal tissues. By leveraging large-scale integration of public RNA-seq datasets spanning 33 tumor types, CIRI-hub provides comprehensive analysis and visualization functions for pan-cancer circRNA analysis. In particular, CIRI-hub permits users to specific circRNAs of their interests or upload their dataset and provides interactively customization of analysis algorithms and plotting parameters on the web interface. Thus, CIRI-hub can serve as an important analysis platform for the effective mining of circRNAs in cancer samples.

## Results

CIRI-hub integrates a compendium of 2187 tumor and 680 normal RNA-seq libraries spanning 33 tissue from previously published resources [18, 20, 22]. To reduce the batch effect of experimental enrichment of circRNAs in different samples and enable the comparative analysis of quantification results, only total RNA libraries without RNase R treatment were collected in our study. Then, the GRCh38 reference genome and the GENCODE v37 annotation were downloaded from the GENCODE [23] website. CircRNAs were subsequently identified using the CIRI2 [24, 25] and CIRIquant [26] pipeline, and their expression level was measured using the counts per million (CPM) method. In sum, a total of 3,176,731 and 580,718 circRNAs were identified in normal and tumor samples, respectively. The gene expression level was estimated using the Hisat2 [27] and StringTie [28] pipeline. Taken together, these collected datasets provide a comprehensive resource for exploring circRNAs in tumors and their matching normal tissues.

To construct a useful resource for most researchers, CIRI-hub included comprehensive analysis results for the collected datasets (**Fig. 1**). Firstly, all circRNAs were annotated into intergenic, intronic, and exonic (exon, 3’UTR, and 5’UTR) types according to their back-splicing junction (BSJ) position. To characterize the differentially expressed circRNAs in each tumor type, the expression profile of each circRNA and their host genes were extracted, and differential expression analysis of both circRNAs and genes between normal and cancer samples was performed in each tissue, respectively. Furthermore, the multiple conservation score (MCS) for circRNAs was calculated using samples from 6 organisms in the circAtlas v2.0 database, which represents the conservation of circRNAs on species, tissues, and individual levels simultaneously. In addition, the sequence features including RBP / miRNA binding site, open reading frames (ORFs), and internal ribosome entry site (IRES) were also predicted, providing important evidence for functional prioritization of identified circRNAs.

**Fig. 1.**
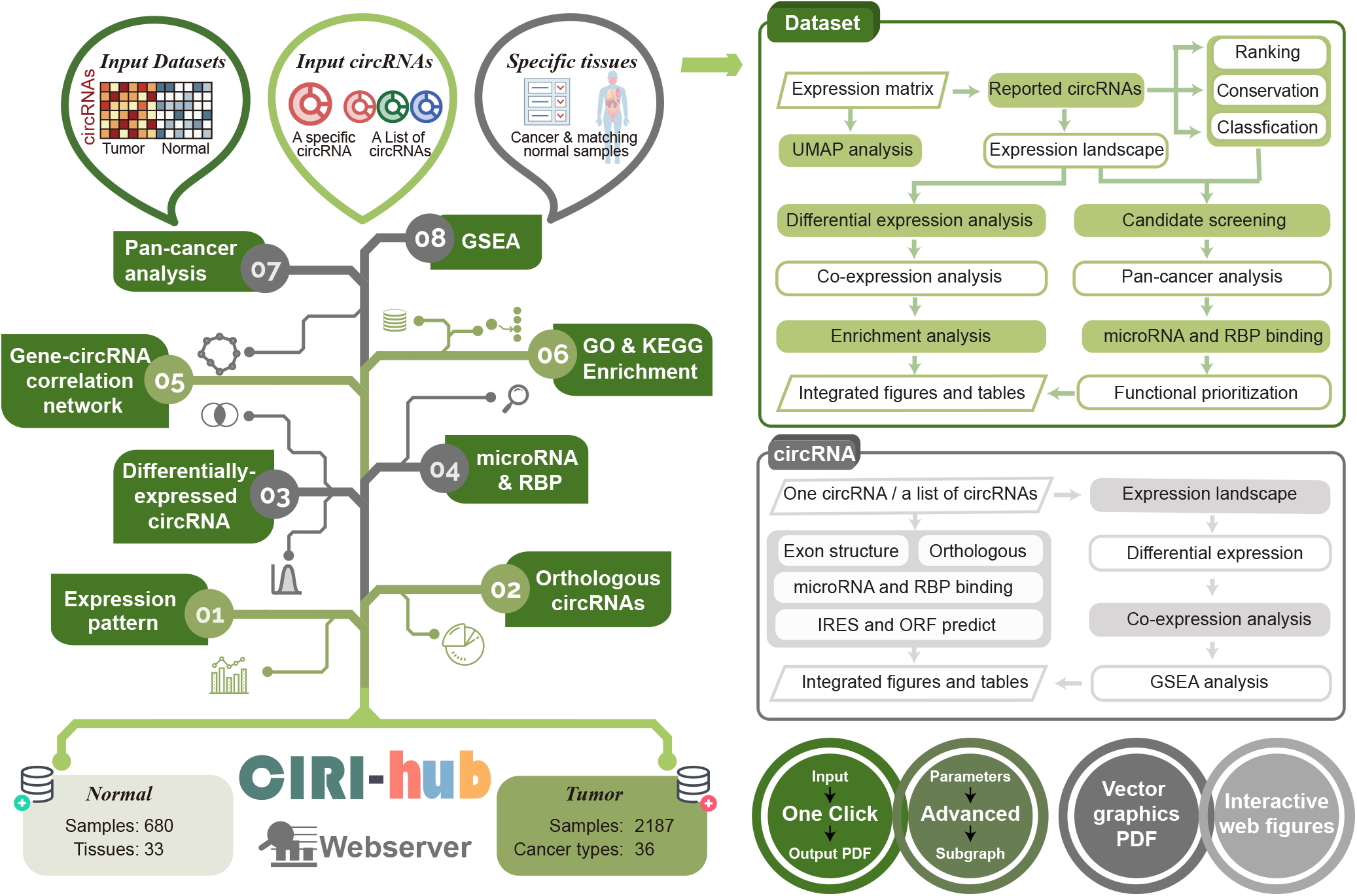
The schematic diagram of the construction of the CIRI-hub webserver.

Based on these collected data and analysis results, CIRI-hub provides one-stop analysis modules for biomarker identification and function prioritization of circRNAs in tumor samples and accepts multiple forms of inputs, including one specific circRNA, a list of circRNAs, and the expression matrix of users’ inhouse RNA-seq datasets. As shown in **Fig.2A**, users can use the “One-click” page to submit their data, and CIRI-hub will return the assigned job id and perform automated analysis and result visualization. Users can either download the output vector graphics in pdf format or customize the plotting parameters (e.g. analysis algorithms, filtering thresholds, color palettes) interactively for each subgraph in the “Browse Details” page. Moreover, the “Advanced” module also implements independent analysis functions including reported circRNAs, type annotation, differential expression, correlation network, conservation score, expression landscape, Gene Ontology & KEGG enrichment analysis, and GSEA analysis (**Fig. 2B**). CIRI-hub also permits users to select interested tumor types and perform these tasks in a tumor-specific context. The pre-analyzed results and integrated figures for each tumor type were also provided on the website.

**Fig. 2.**
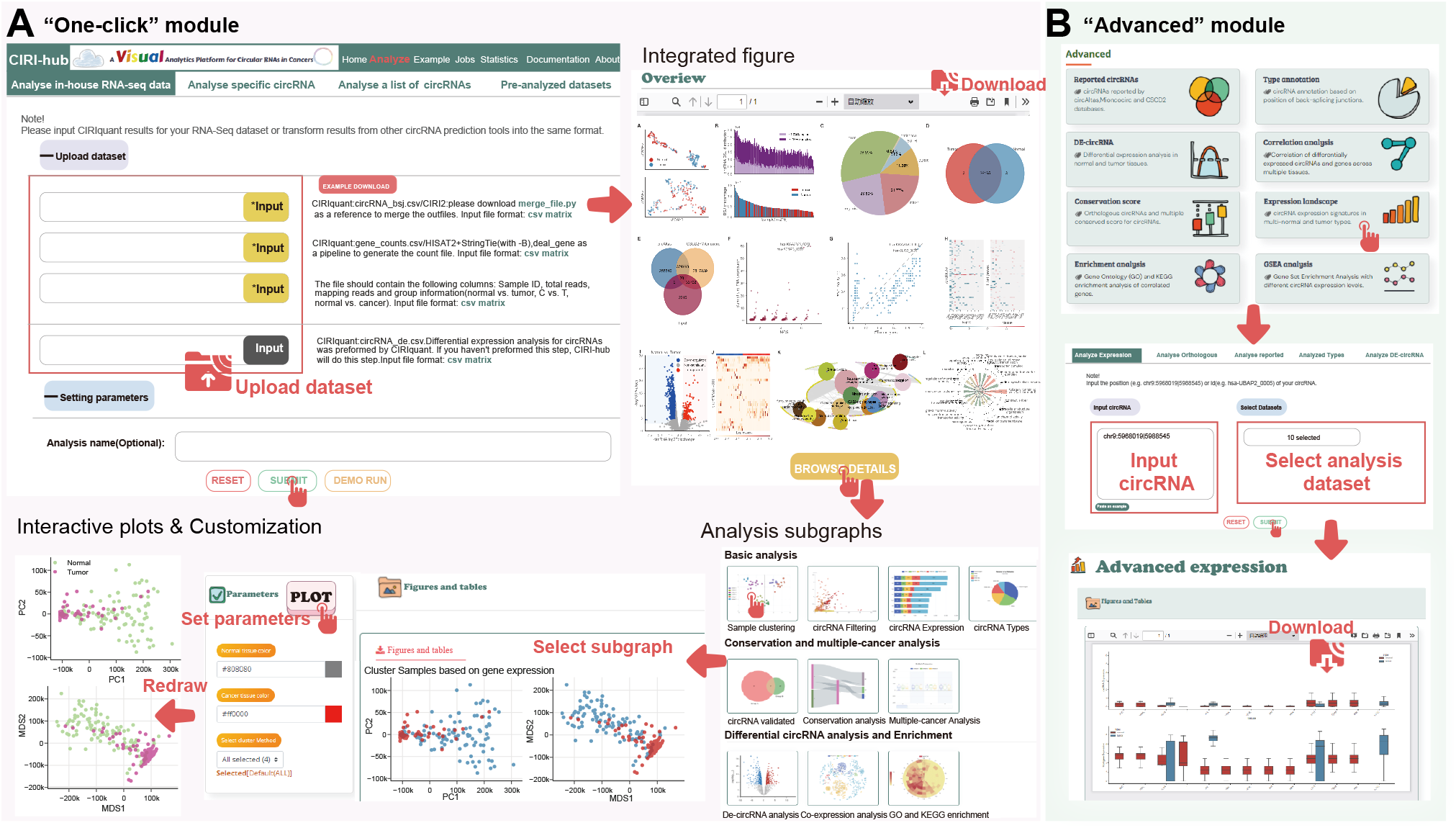
The functions of the CIRI-hub webserver. **A**. “One-click” module for automated analysis. **B**. “Advanced” module for performing analysis functions separately.

For users that have in-house RNA-seq datasets, CIRI-hub can serve as a comprehensive analysis and visualization platform. Users can generate the gene and circRNA expression matrix using CIRIquant with their RNA-seq data, then upload to the “One-click” page of CIRI-hub for automated analysis. Firstly, the number of detected circRNAs and BSJ reads for each sample are summarized, and CIRI-hub offers four different dimension reduction analysis algorithms (PCA, MDS, t-SNE, and UMAP) to examine whether the normal and tumoral samples can be clearly distinguished using expression levels of genes or circRNAs (**Fig. 3A**). Then, all detected circRNAs are classified into different types according to the BSJ position. To ensure the reliability of circRNA identification, the number of circRNAs detected in normal and tumor samples are calculated. Next, CIRI-hub implements a pan-cancer analysis of reported circRNAs in 33 tumor types. All detected circRNAs are compared to the collected datasets, and the multiple conservation score and the occurrence frequency of these reported circRNAs are plotted, where conserved circRNAs are explicitly marked in the diagrams. The expression change of these circRNAs and their host genes in various tumor types are also plotted, which allows the users to select highly conserved circRNAs or novel tumor markers. Meanwhile, CIRI-hub also performs differential expression analysis using the input datasets. The log2 fold change and q-value of circRNAs are calculated using CIRIquant and plotted in the volcano plot. Besides, CIRI-hub also provides the correlation of expression changes of circRNAs and their host genes and performs Gene Ontology, KEGG, and GSEA enrichment analysis of differentially expressed circRNAs (**Fig. 3B**).

**Fig. 3.**
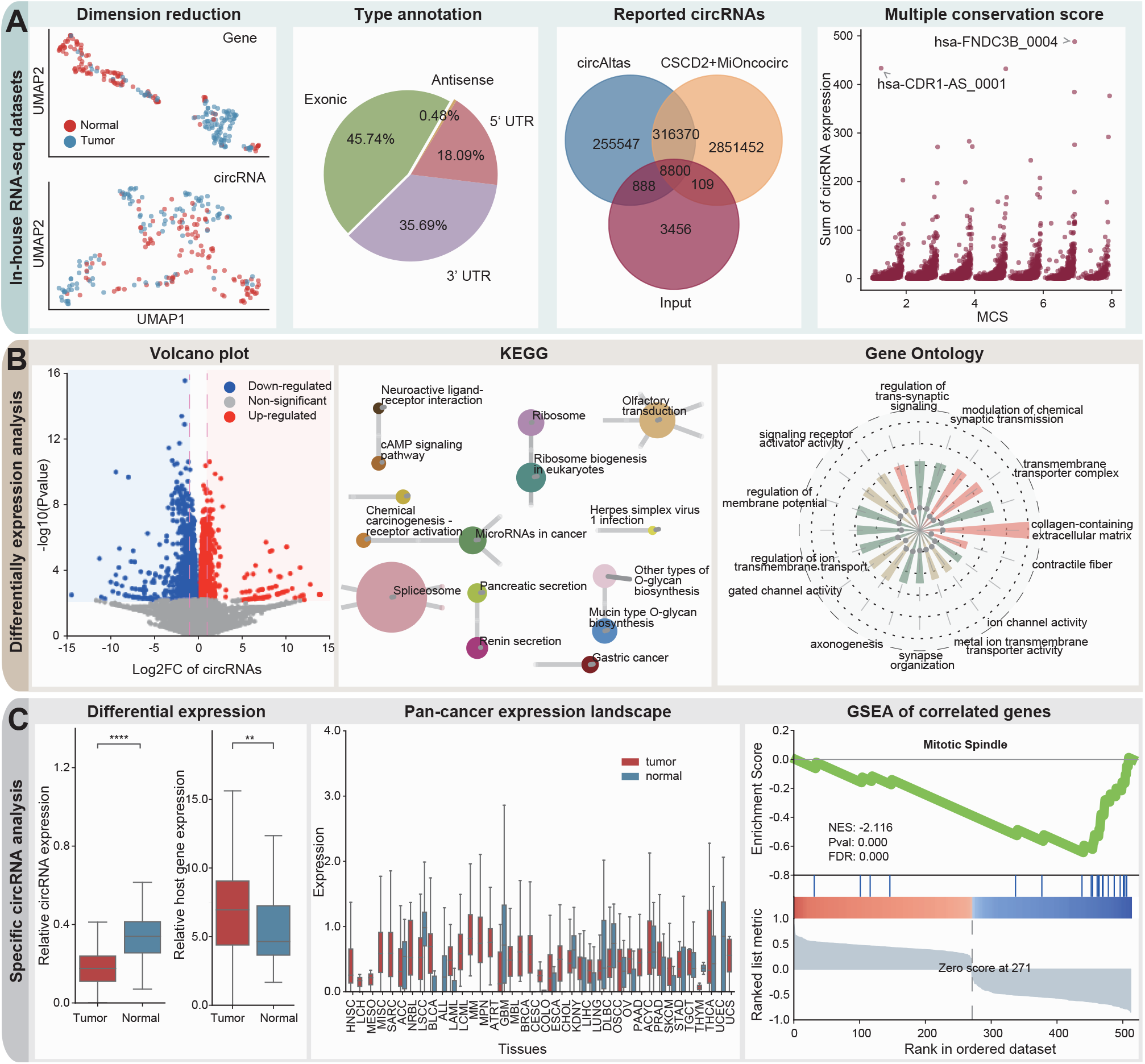
Example output of the CIRI-hub webserver. **A**. Example output figures for analyzing in-house RNA-seq datasets. **B**. Example output figures for differential expression and functional enrichment analysis. **C**. Example output of analysis for specific circRNA.

Next, users can download the differential expression results, and use a list of candidate circRNAs for downstream analyses. The “Analyze a list of circRNAs” page provides detailed information, including MCS score, change of circRNAs, and the host genes expression in each tumor type, which provides essential information for screening potential biomarkers in different tumor types. For specific circRNA, the “Analyze specific circRNA” module provides its expression landscape across normal samples of six organisms and tumor samples spanning 33 tumor types. Moreover, CIRI-hub can find genes that are highly correlated with the input circRNA in the co-expression network and perform GSEA for functional annotation (**Fig. 3C)**. Besides, the sequence features including RBP / miRNA binding site, coding potential, and IRES are also plotted in the combined figures, providing comprehensive evidence for selecting circRNA candidates for further experimental validation.

CIRI-hub is implemented in the high-performance computing cluster of Beijing Institutes of Life Science, Chinese Academy of Sciences. The webserver jobs are processed on a single node with 64 CPU cores and 512 GB RAM. All data are stored in 3.0 PB of high-performance storage, which ensures the high availability of this webserver. All analysis and visualization results can be obtained from a temporary web link that will be stored for 7 days.

## Conclusion

In conclusion, CIRI-hub for the first time provides a user-friendly platform for one-stop analysis and visualization of the users’ circRNA datasets. By integrating a large compendium of RNA-seq datasets, CIRI-hub provides comprehensive analysis functions of circRNAs in tumors and matching normal tissues. CIRI-hub consists of three major components: (1) “Analyze in-house RNA-seq data” module for analyzing the expression change of circRNAs in the users’ dataset and collected datasets; (2) “Analyze a list of circRNA” module for filtering highly conserved and novel circRNA biomarkers from the candidate list; (3) “Analyze specific circRNA” module to provide insights into the possible biological function of certain circRNA in multiple tumor types. CIRI-hub also integrates multiple analysis functions and users can easily access pre-analyzed results for each tumor type. All these analysis results are assembled into publication-quality graphics that can be interactively customized on the web interface. Thus, we believe the CIRI-hub will facilitate more efficient utilization of public RNA-seq resources and can serve as a useful web server for mining circRNAs in tumor and normal samples.

## Abbreviations

KEGG: Kyoto Encyclopedia of Genes and Genomes
GSEA: Gene Set Enrichment Analysis

## Declarations

### Ethics approval and consent to participate

Not applicable

## Consent for publication

Not applicable

## Availability of data and materials

The CIRI-hub web server can be accessed at http://159.226.67.237/zhao/Database/CIRIhub/index.html. The RNA-seq datasets used to construct the webserver ca n be accessed from the website.

## Authors’ contributions

F.Z. conceived the project. W.W. and J.Z. analyzed the data. W.W. designed the webserver. J.Z., W.W., and F.Z. wrote the manuscript. The authors read and approved the final manuscript.

## Acknowledgments

Not applicable

